# Resolving single-cell heterogeneity from hundreds of thousands of cells through sequential hybrid clustering and NMF

**DOI:** 10.1101/608869

**Authors:** Meenakshi Venkatasubramanian, Kashish Chetal, Gowtham Atluri, Nathan Salomonis

**Author notes:** **Corresponding Author:** Nathan Salomonis, Ph.D, Division of Biomedical Informatics, Cincinnati Children’s Hospital Medical Center, 3333 Burnet Avenue, Cincinnati, OH, 45229, USA.

## Abstract

The rapid proliferation of single-cell RNA-Sequencing (scRNA-Seq) technologies has spurred the development of diverse computational approaches to detect transcriptionally coherent populations. While the complexity of the algorithms for detecting heterogeneity have increased, most existing algorithms require significant user-tuning, are heavily reliant on dimensionality reduction techniques and are not scalable to ultra-large datasets. We previously described a multi-step algorithm, Iterative Clustering and Guide-gene selection (ICGS), which applies intra-gene correlation and hybrid clustering to uniquely resolve novel transcriptionally coherent cell populations from an intuitive graphical user interface. Here, we describe a new iteration of ICGS that outperforms state-of-the-art scRNA-Seq detection workflows when applied to well-established benchmarks. This approach combines multiple complementary subtype detection methods (HOPACH, sparse-NMF, cluster “fitness”, SVM) to resolve rare and common cell-states, while minimizing differences due to donor or batch effects. Using data from the Human Cell Atlas, we show that the PageRank algorithm effectively down samples ultra-large scRNA-Seq datasets, without losing extremely rare or transcriptionally similar distinct cell-types and while recovering novel transcriptionally unique cell populations. We believe this new approach holds tremendous promise in reproducibly resolving hidden cell populations in complex datasets.

**Highlights:**

- ICGS2 outperforms alternative approaches in small and ultra-large benchmark datasets
- Integrates multiple solutions for cell-type detection with supervised refinement
- Scales effectively to resolve rare cell-states from ultra-large datasets using PageRank sampling with a low memory footprint
- Integrated into AltAnalyze to enable sophisticated and automated downstream analysis

## INTRODUCTION

Recent advances in single cell RNA sequencing (scRNA-Seq) provide exciting new opportunities to understand cellular and molecular diversity in healthy tissues and disease. With the rapid growth in scRNA-Seq, numerous computational applications have been developed that address diverse technical challenges such as measurement noise/accuracy, data sparsity and high dimensionality to identify cell heterogeneity within potentially complex cell populations. Most applications consist of a shared set of steps, including: 1) gene filtering, 2) expression normalization, 3) dimensionality reduction, and 4) clustering ^1^. While the specific algorithms and options used for these steps varies significantly among applications, most approach rely heavily on dimensionality reduction techniques, such as PCA, t-SNE and UMAP to select features and define cell populations. As noted by others ^1^, the reliance on such techniques has several limitations, including insensitivity to non-linear separation of the components (PCA), loss of global structure due to a focus on local information (t-SNE) ^2^ and inability to scale to high-dimensions (UMAP) ^3^, resulting in a significant loss of information during projection.

While a number of methods exist to identify clusters from large lower dimensional projections, including DBSCAN, K-means, affinity propagation, Louvain clustering and spectral clustering, these, as well as other approaches require proper hyperparameter tuning. Identifying these parameters is non-intuitive and often requires multiple rounds of analysis. To address this concern, consensus-based approaches that considers the results from multiple runs with different parameters have been developed, such as SC3 ^4^, however the ultimate selection of the parameters remains user dependent and is not automated. Thus, identifying the right approach and parameters for dimension reduction or clustering for each new dataset remains time consuming and technically challenging. The increasing production of atlas sized datasets highlights the important need for highly scalable and automated computational approaches that can rapidly identify common and extremely rare populations with minimal user parameter tweaking ^5^.

Here, we present the next iteration of our previously described approach Iterative Clustering and Guide-gene Selection (ICGS) ^6^. Unlike alternative approaches, ICGS iteratively identifies core variable gene expression programs from a data matrix through multiple rounds of hybrid clustering (HOPACH ^7^), selection of maximally informative guide-genes and expression correlation. ICGS2 incorporates multiple additional downstream methods to improve subtype detection and reduce noise, including sparse-NMF ^8^, cluster fitness, SVM ^9^ and automated methods to improve parameter estimation. Here, we introduce a novel intelligent sampling-based strategy for extremely large datasets to capture the most informative cells for downstream unsupervised analyses. ICGS2 is an easy-to-use automated pipeline that requires minimal user inputs and no a priori details regarding the cells or the genes analyzed. Through integration with AltAnalyze ^10^, the workflow can be run from the command-line or an intuitive graphical user-interface and includes a large repertoire of user-friendly integrated downstream analysis tools (e.g., cell-type prediction, differential expression, pathway analysis). We demonstrate improved performance of ICGS2 when compared to diverse alternative algorithms applied to scRNA-Seq datasets of varying size and complexity. Importantly, this approach remains scalable to ultra-large data sets, without sacrificing sensitivity. These evaluations demonstrate that ICGS2 represents an automated, scalable, accurate and easy-to-use platform for next-generation scRNA-Seq analysis.

## METHODS

### Algorithm Design

The software ICGS2 has been developed as part of an open-source python toolkit, AltAnalyze, with a complete documentation of its use, algorithms and optional user-defined parameters (https://altanalyze.readthedocs.io/en/latest/). ICGS2 identifies cell clusters through a five-step process: 1) PageRank-Down-sampling (optional), 2) feature selection-ICGS2, 3) dimension reduction and clustering (sparse-NMF), 4) cluster refinement (MarkerFinder algorithm), and 5) cluster re-assignments (SVM) (Fig. 1a). AltAnalyze includes support for multiple input format including: A) an already normalized expression file, such as counts-normalized, non-log or log2, with genes as rows and samples as columns, B) 10× Genomics (version 1.0-3.0) produced filtered sparse matrix results (.mtx, HDF5), C) genome-aligned BAM files, or D) FASTQ files for individual cells. A tab-delimited gene-counts matrix can be normalized prior to import using the module CountsNormalize. The principle steps of this program are:

**Figure 1.**
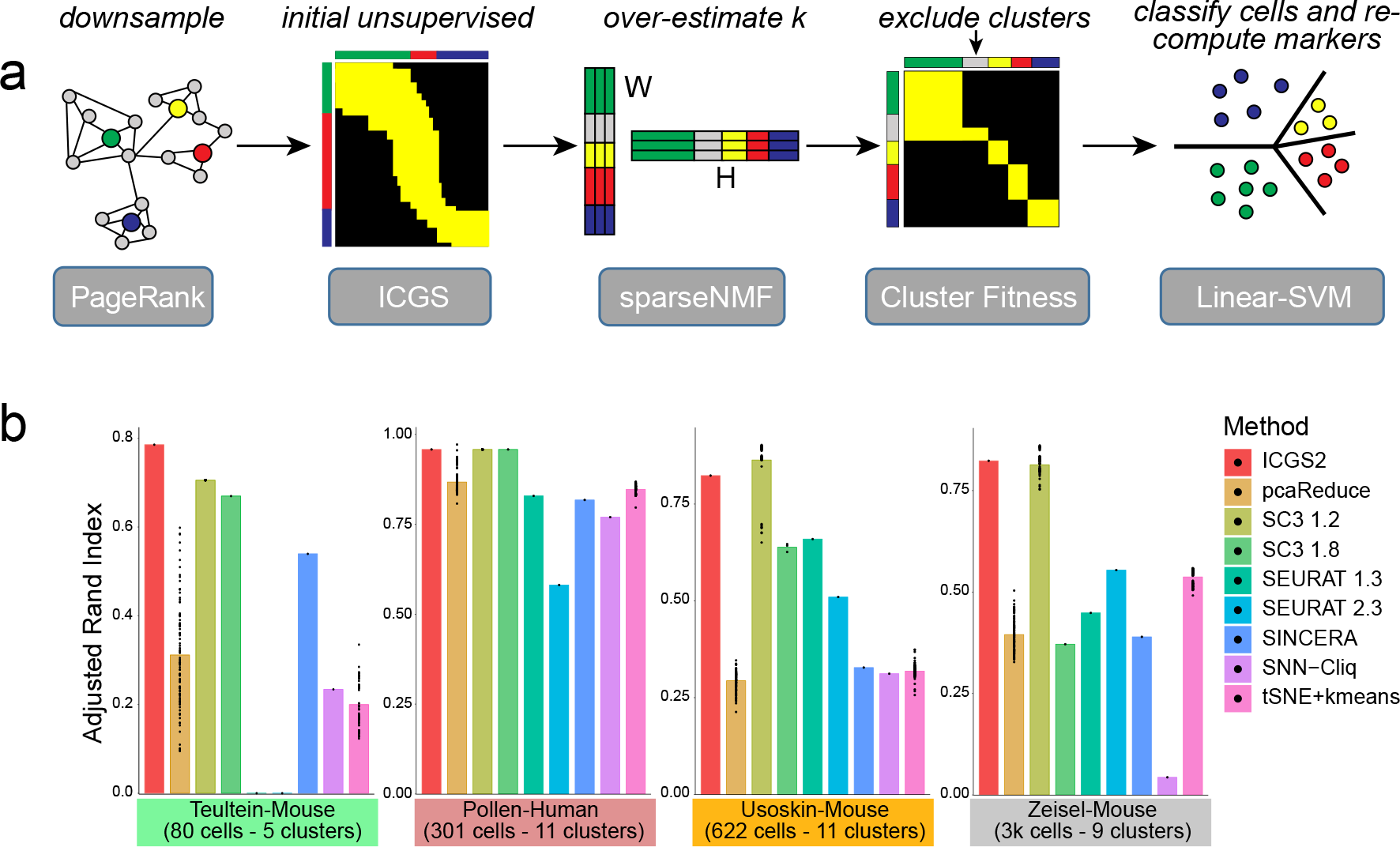
Performance of ICGS2 against diverse alternative unsupervised scRNA-Seq algorithms. a) Overview of the ICGS2 workflow for single cell RNA-Seq population prediction. These steps include: 1) PageRank-Down-sampling (optional), 2) feature selection (ICGS), 3) dimension reduction (Sparse-NMF), 4) cluster refinement/exclusion (“fitness”) and 5) cluster assignments (Linear-SVM). b) Comparison of ICGS2 to previously evaluated algorithms to detect prior defined cell populations from distinct “silver” standard datasets of varying size and complexity using the Adjusted Rand Index (ARI). Specific algorithms evaluated, with specific versions of these tools indicated.

#### Step 1: PageRank-Down-sampling (recommended for datasets with > 2,500 cells or user-defined)

a. <u>Selection of variable genes</u>: ICGS2 imports an input expression file processed from AltAnalyze (automatically log2 normalized, protein-coding genes and initial ICGS variance filtered) and identifies the top 500 genes with the highest dispersion. Dispersion for each gene is calculated as the ratio of the variance divided by its mean. The resulting PageRank input file is filtered for these genes.
b. <u>Graph construction</u>: Since the filtered input is not a graph, a graph representation of the dataset is created by finding the k-nearest neighbors (k =10 default) for each cell. Identification of the k-nearest neighbors is performed using efficient python package called Annoy ^11^ and the graph is created using the networkx python package. (Optional) For very large scRNA-Seq datasets (n>25,000), an initial down-sampling is performed using a community analysis approach, Louvain clustering (community python library). Louvain clustering is performed with the lowest possible resolution (r=0) to find maximal clusters (smallest communities). This value indicates at which level to cut the clustering dendrogram, with 0 resulting in the most granular and −1, the least. Once each community is identified using Louvain clustering, it is then sampled to identify medoid cells that are representative of the community. A medoid for a community is defined as

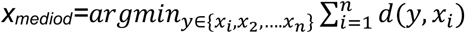

where *x*_*i*_, *x*_2_, …. *x*_*n*_ are the cells of a community, n is the total number of cells in the community, and d is the distance function. The distance function considered here is euclidean distance (sklearn.metrics.pairwise). The total number of medoids sampled for a community is given as:

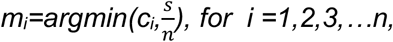

where n is the total number of communities, c_i_ is the total number of cells in the community i, m_i_ is the number of medoids to be identified for the community i and s is the required down-sampling number (default=10,000). This step is performed to increase the speed and efficiency in processing large datasets and reduce the sample space (cells) for PageRank (step 3). Steps a and b are then repeated on the Louvain sampled data.
c. <u>PageRank</u>: Once the graph is generated, a score is calculated for each cell based on PageRank score (networkx python library). The top 2,500 cells (by default) with the highest PageRank score is selected and used for the remaining analysis. For datasets of potentially millions of cells, these default thresholds should likely be increased to accommodate potentially hundreds of cell types.

#### Step 2: Feature selection

While feature selection in ICGS2 is the same as in the original version ICGS, the associated thresholds are automatically determined, including the correlation cutoff appropriate for the dataset. In brief, ICGS identifies correlated gene modules through pairwise correlations of variable genes (Pearson>user supplied threshold), followed by multiple rounds of HOPACH clustering of genes and cells and determination of representative marker genes (guide-genes) for supervised correlation analysis. Guide-gene selection enables the direct exclusion of cell-cycle gene expression modules by exclusion of guide-associated cell cycle genes prior to supervised correlation of those guides. ICGS2 begins with a default Pearson rho>0.2 for the identification of correlated genes, however, if the number of initial correlated genes is > 5,000, the cutoff is automatically incremented by 0.1 and the correlation step is re-iterated until this cutoff is met. By default, only protein-coding genes are considered with exclusion of mitochondrial genome, L ribosomal and S ribosomal genes. 10× Genomics data is automatically imported and normalized (counts per gene divided by the total counts per barcode multiplied by a 10,000).

#### Step 3: Dimension Reduction with sparse-NMF (SNMF)

To improve the delineation of cell-clusters following HOPACH clustering in ICGS, SNMF is applied to the clustered cells to refine population detection. SNMF uses a L1-norm minimization and is solved using a fast nonnegativity constrained least squares algorithm (FCNNLS)^8^. The Guide3 results from ICGS are produced as previously described ^6^. To estimate the number of ranks (i.e., clusters) for SNMF, the ICGS Guide3 (final correlation heatmap text output of ICGS) matrix is z-score normalized and its eigenvalues are calculated. The number of clusters is estimated as 2\**k*, where k is determined by the number of eigenvalues that are significantly different with *P* < 0.001 from the Tracy–Widom distribution ^4^whose mean is 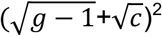 and standard deviation is:

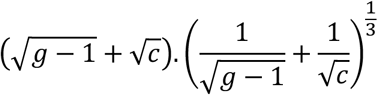

where is the g is the number of genes and c is the number of cells ^4^.

Dimension reduction is performed on the ICGS Guide3 results using SNMF/R which is available in the ‘nimfa’ python package. Given an input matrix *c × g* where c is the number of cells and g is the number of genes, the SNMF/R factorization returns two sub-matrices called the basis matrix, W with the dimensions *c × r*, where c is the number of cells and r is the number of ranks and the coefficient H matrix with the dimensions *g × r*, where g is the number of genes and r is the number of ranks. For each cell, its assignment was based on its largest contribution in W. All the parameters are set to default as per the package except the rank.

#### Step 4 Marker gene selection (cluster fitness)

In some cases, the clusters identified in step 3 will be weakly defined by unique gene expression. To identify rigorously defined cell-clusters with unique gene expression for downstream cell-cluster assignment (all-cells, not just down-sampled), ICGS2 applies the MarkerFinder algorithm, which is a component of AltAnalyze ^6^. MarkerFinder identifies genes that are positively correlated with an idealized cluster-specific expression profile (1 or 0). For each SNMF cluster, a reference is created where cells belonging to the group are assigned 1 and the remaining cells are assigned 0. Each gene is correlated to the references and assigned to a particular cluster based its highest Pearson correlation (rho). Using the initial correlation cutoff identified for ICGS pipeline, SNMF cell clusters with less than 2 genes above the supplied rho threshold are excluded from the analysis. As such, centroids will be derived for only clusters with unique gene expression for supervised assignment to those final cluster. The top 60 Pearson correlated genes for each SNMF cluster with a rho>0.3 are considered for the remaining SNMF groups. As such, this method addresses the vital unmet need to exclude clusters that specifically result from noise or low-depth sequencing (assuming transitional or mixed-lineage states are associated with some uniquely expressed genes).

#### Step 5: Cell cluster assignment (linear SVC)

Using the marker genes identified for sufficiently fit clusters, cluster medoids are determined based on the cells assigned to the specific SNMF clusters. Next, a Linear SVM model with a linear kernel is constructed. This model is applied to all the cells in the dataset and re-classified based on the training models. ICGS2 uses the LinearSVC option in scikit-learn (default parameters).

### User Parameters

By default, ICGS2 includes built-in automated parameter estimation for its correlation cutoff (ICGS and MarkerFinder), estimation of cluster number (rank estimation for sparse-NMF) and number of cells to down-sample. These defaults can be explicitly set by the user to force the software to identify more or less clusters/heterogeneity. Additionally, ICGS has default options which can be modified by the user including: 1) number of variable cells, 2) gene expression fold-cutoff, 3) protein-coding gene filter, 4) exclusion of cell-cycle effects and 5) HOPACH clustering metric (Correlation, Euclidean, Cosine). For evaluation of these methods, the software defaults have been used.

### Benchmarking

To evaluate the performance of ICGS2, we compare cell clustering results to known reference clusters identified by previous authors of the different datasets tested. Adjusted RAND Index ^12^ which has been used previously for these validations have been used to test the tools rigorously.

### Sample Datasets

Sample datasets that can be tested with ICGS2 are provided with AltAnalyze software. This include a sample 10× Genomics dataset (sparse matrix files) and Fluidigm scRNA-Seq data. Please see the folder DemoData and associated information and instructions for analysis in the AltAnalyze program folder. A YouTube tutorial for running ICGS2 with this data (along with downstream comparison analyses using the tool cellHarmony) can be found at: https://youtu.be/mRiGIz-zV80.

### Software Availability

ICGS2 is a part of AltAnalyze, starting with version 2.1.2. AltAnalyze is available as open-source python code (https://github.com/nsalomonis/altanalyze), PyPI installation (pip install altanalyze) and as pre-compiled binaries with dependent libraries included (http://www.altanalyze.org). Through AltAnalyze, this software can be run through an intuitive graphical user interface (https://altanalyze.readthedocs.io/en/latest/RunningAltAnalyze/) or using command-line options in the various implementations (https://altanalyze.readthedocs.io/en/latest/CommandLineMode/). Online video tutorials are provided for example input datasets (http://www.altanalyze.org).

### Software Outputs

Primary ICGS2 results include marker gene heatmaps with likely predicted cell types (down-sampled and all cells), UMAP projection, marker genes associated with each cell population and ranks (text file), SVM scores (text file) and cell-to-cluster (text file) associations within the ICGS-NMF and NMF-SVM folders. Secondary results include differential expression results between clusters, protein-protein and protein-DNA predicted interactions among these genes (network plots), QC plots, LineageProfiler cell-type predictions and GO-Elite pathway/Ontology/gene set enrichments by default.

## RESULTS

To improve the prediction of discrete cell states from diverse possible single-cell RNA-Seq datasets, we developed a significantly improved iteration of our software ICGS. These new methods were built on-top of ICGS rather than creating a new method from scratch, as this software has several potential fundamental advantages over existing approaches. These advantages include ease-of-use (graphical and non-graphical user interfaces), a lack of reliance on dimensionality reduction to identify initial cellular and gene expression heterogeneity (guide-gene based discovery), automated data visualization outputs (heatmap, UMAP), methods for cell-type prediction and embedded pathway/network analyses. We previously demonstrated that use of this approach improves the delineation of rare transcriptionally distinct populations while minimizing “batch” or donor-bias through the selection of highly coherent gene expression clusters derived through intra-correlation of genes ^6, 13^. To improve the delineation of rare transcriptional states, we have augmented the core ICGS algorithms with rigorous methods for determining biologically valid clusters (SNMF, SVC, cluster fitness), automated cluster number determination, introduced a new method for accurate down-sampling (e.g., PageRank) for large scRNA-Seq datasets and added new methods for data visualization (UMAP) (Fig. 1a). These methods were designed to increase sensitivity of ICGS to identify important rare cell populations in datasets with potentially hundreds of thousands of cells.

### ICGS2 outperforms alternative algorithms for established benchmarks

ICGS2 was tested against multiple gold- and silver-standard datasets, with varying technologies, dimensionality and complexity. To benchmark the core subtype identification module, four datasets previously used for benchmarking the SC3 algorithm against diverse established algorithms. The datasets Zeisel ^14^, Pollen^15^, Usoskin ^16^ and Treutlein ^17^ were selected particularly for their size and number of clusters. Further, these datasets had variable performance for the different methods compared. For comparison with ICGS2, we consider the previously benchmarked tools SC3 v.1.2 ^4^, SEURAT v.1.3 ^14^, SINCERA ^18^, SNN-Cliq ^19^, tSNE+kmeans, as well as updated versions of SC3 (version 1.8) and SEURAT (version 2.3) ^20^. The Adjusted Rand Index (ARI) was used for the evaluation metric and author provided labels were used as ground state truth. Incorporating the original and our ARI measurements, we find that ICGS2 has improved or equivalent performance to all other methods tested, including SC3 v.1.2 (Fig. 1b). As the newer version of SC3 had decreased performance over the original tested, we reran the data, for SC3 v.1.2 and Seurat v.1.3 and validated the previously reported ARIs. These results indicate that ICGS2 collectively outperforms other methods on datasets of distinct size and complexity.

### Identification of distinct hematopoietic subtypes in the Human Cell Atlas

We recently performed a comprehensive analysis of eight independent donor bone marrow scRNA-Seq samples collected and profiled from Human Cell Atlas (HCA) initiative ^21, 22^. This analysis defined 35 distinct hematopoietic cell populations from over 100,000 cells. Although the workflow applied from this analysis relied on ICGS, ICGS was run independently on the cells from each eight donor individually, prior to those results being aggregated and used as references for cell alignment using a novel classification strategy. This analysis produced both a combined dataset with all mature and progenitor cells and a separate analysis in which selectively defined refined populations in presumptive bone marrow progenitors (11,548 CD34+ cell clusters). As these populations were independently verified using prior sorted-population transcriptomic references and/or marker genes and were found to largely exclude donor-driven effects ^21^, we consider these predictions as additional “silver” standards. When applied to this smaller dataset of progenitors (11,548 cells), ICGS2 had the greatest relative performance (ARI), relative to the algorithms SC3 v.1.8, Seurat v.2.3, Scanpy ^23^ and two recent versions of the Monocle software (version 2 ^24^ and version 3 ^5^) (Fig. 2a).

**Figure 2.**
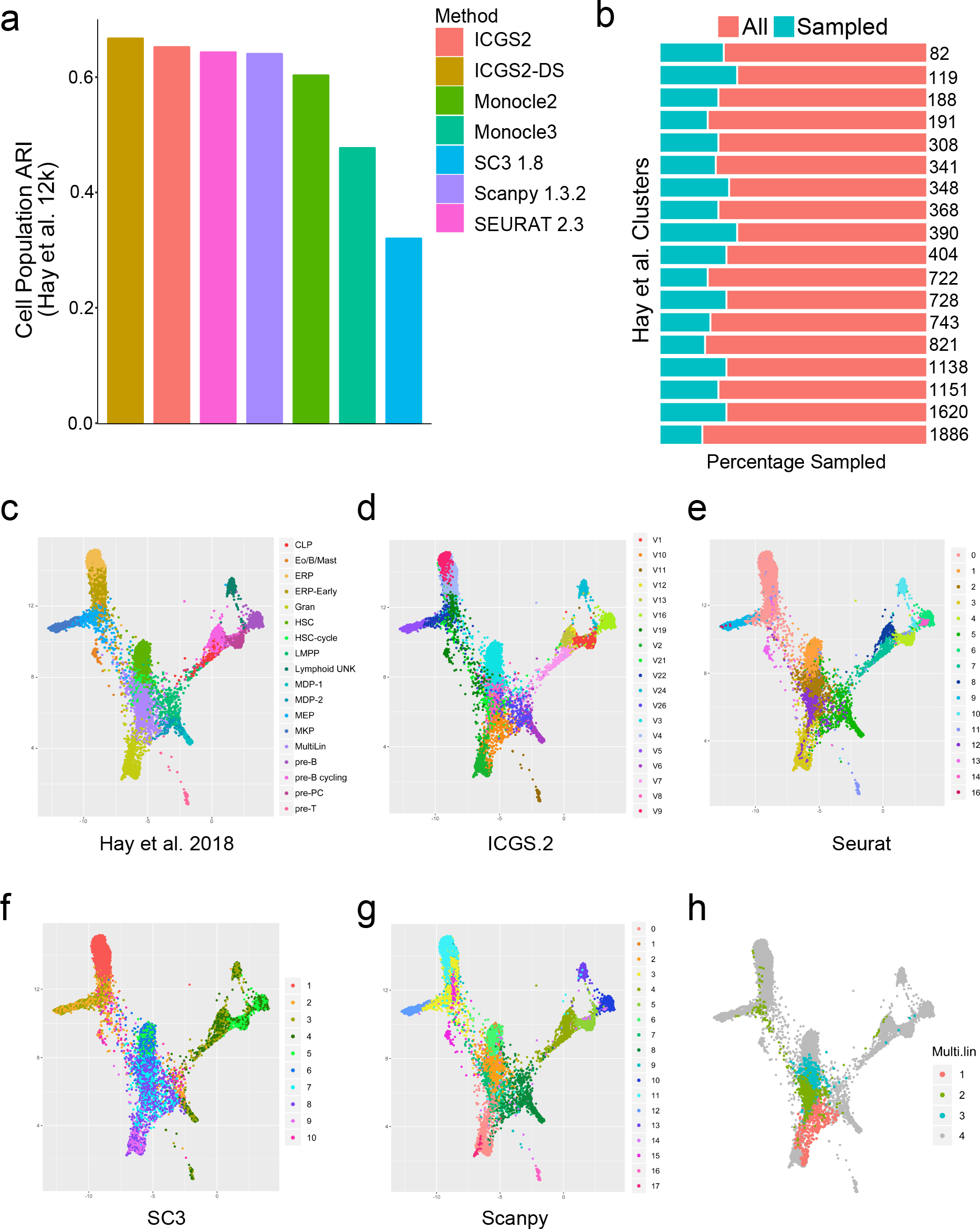
Delineation of discrete and transitioning populations in bone marrow progenitors. a) Detection of prior-defined Human Cell Atlas (HCA)(n=11,568) bone marrow progenitor (BMP) clusters using ICGS2 and ICGS2-DS (down-sampled) compared to Seurat 2.3, Scanpy, SC3 1.8 and Monocle 2 and 3. To maximize the ARI score for each approach, if multiple clusters were predicted for a single reference cluster (Hay et al.), these clusters were combined prior to scoring. b) Frequency of PageRank down-sampled cells out of the total number of previously defined BMP clusters (n=18). The total number of cells in each cell population are shown to the right of the plot. Visualization of cell cluster assignments using the software SPRING of: c) prior-defined HCA BMP clusters, d) ICGS2 using down-sampling (ICGS2-DS), e) Seurat v2.3, f) SC3, and g) Scanpy. h) Spring visualization of novel multi-lineage populations identified uniquely by ICGS2.

### Optimized population discovery from large scRNA-Seq datasets

ICGS2 is dependent on HOPACH clustering and SNMF which are computationally expensive with increasing dataset size. As such, it is not immediately applicable to ultra-large datasets, such as the HCA bone marrow compendium, hence, we implemented a new method for intelligent down-sampling of the cellular data, prior to the existing SVM classification of cells. While approaches such as SC3 apply random down-sampling, this procedure is likely to miss rare but biologically important cell populations. Alternatively, a recent down-sampling single-cell method (BigScale) applies k-nearest neighbor approach that is effective at preserving heterogeneity in large scRNA-Seq datasets, but requires defining the number of nearest neighbors a priori ^25^. To address this challenge, ICGS2 applies the Google PageRank algorithm ^26^, a graph-based algorithm, originally designed to identify the most frequently visited websites. This approach prioritizes the selection of interspersed nodes in the larger graph, with minimum representation bias. Using the PageRank score, ICGS2 identifies the top 2,500 representatives cell profiles (by default) for large datasets. To evaluate this down-sampling technique, we again used the progenitor population dataset (HCA, n=11,548). We compared the percentage of cells retained for each known group and found that the down-sampling approach consistently identified around 17-26% from the total 22% cells considered (Fig. 2b). The results of the original ICGS2 and down-sampled ICGS2 were highly concordant, with an ARI of 86%. We attributed the differences in the ARI to the slight differences in the assignment of cells in particular highly related transitional states. While, none of the evaluated scRNA-Seq algorithms were able to replicate some of the original author transcriptionally distinct clusters (two separate Monocytic Dendritic Precursor (MDP) populations, Hematopoietic Stem Cell (HSC) in cycle versus HSC), both down-sampled and complete ICGS2 selectively identified Common Lymphoid Progenitors (CLP) and Lymphoid-primed multipotent progenitors (LMPP) not identified by the other algorithms. ICGS2 further found additional granularity in the original annotated presumptive Multi-Lineage progenitor (Multi-Lin) cells (Fig. 2c-g). This delineation is supported by unique gene expression present in these additional Multi-Lin subsets, with skewed granulocytic, MDP or HSC gene expression. Visualization of these subsets in the software SPRING ordered these three Multi-Lin populations groups along separate connected trajectories further supporting the validity of these predictions (Fig. 2h).

### ICGS2 uniquely identifies novel subpopulations in ultra large datasets with minimal donor effects

We next compared the performance of ICGS2 in the complete HCA dataset against other approaches compatible with ultra-large scRNA-Seq datasets. For datasets of >25,000, Louvain clustering is performed with minimum resolution to down-sample the data to around 10,000 cells and construct the graph, PageRank then is applied to identify the top 2,500 representative cells. This step is optional to improve the performance of graph construction for the sampling. As with the CD34+ restricted dataset, we compared the percentage of the cells sampled for each of the author provided labels for the 2% of cells sampled by this procedure. On average, 10% of cells in populations with less than 1,000 cells were selected from by this method and 3% for populations with more than a thousand cells. At least 6 representative cells per cell cluster were selected by this down-sampling method for all 35 previously defined bone marrow cell populations (Fig. 3a). To compare its ability to detect cell populations, ICGS2 was again evaluated relative to Seurat (v2.3), Monocle3 and Scanpy, which have previously shown to effectively handle large datasets. While runtime on this dataset ranged from 15 minutes (Scanpy) to 6 hours (Seurat), ICGS2 proved to be the most memory efficient method, while remaining relative fast (2 hours) (Table 1). We attempted to run SC3, however, this approach reached its memory limit with 256GB of RAM (estimate k-step). BigScale was excluded from evaluation as it is currently compatible only with Windows operating systems (Matlab license required). To assess the contribution of donor driven effects in the clusters obtained, Seurat was also run using the canonical correlation analysis option (Multi-CCA) with the different donors considered as different datasets. When comparing these different methods, ICGS2 was found to have the greatest ARI for the 35 prior assigned cell-populations (Fig. 3b). In addition, as we predicted, ICGS2 identified clusters were the least confounded by donor effects, including those identified by Seurat Multi-CCA (Fig. 3c). In addition to the previously described bone marrow cell populations, ICGS2 uniquely identified distinctive additional subtypes of T-cells, Erythroblasts and Dendritic cells (DC) which were not previously identified nor identified by the other approaches. For example, each DC cell cluster was found to expresses unique marker genes with established roles in functionally distinct DC subsets (plasmacytoid, maturing CD1c+, CD1c+ and CD8+) (Fig. 3f-i) ^27, 28, 29, 30, 31^. It is important to note that all approaches tested failed to sufficiently define all of the discrete CD34+ progenitor cell populations in the entire HCA dataset that were clearly resolved from the independent analysis of these cells, suggesting that these methods are far from perfect. Nonetheless, ICGS2 not only outperformed these other approaches, but identifies unique cell populations that align to prior knowledge.

**Table 1:** Computational benchmarking of ICGS2 relative to alternative approaches.

| **Application** | **Maximum**<br>**Memory Usage** | **Processing**<br>**Time** |
| --- | --- | --- |
| ICGS2 | 10.4 GB | 121.2 min |
| Scanpy | 33.9 GB | 14.9 min |
| Monocle3 | 170.4 GB | 81.3 min |
| Seurat | 160.1 GB | 355.1 min |

**Figure 3:**
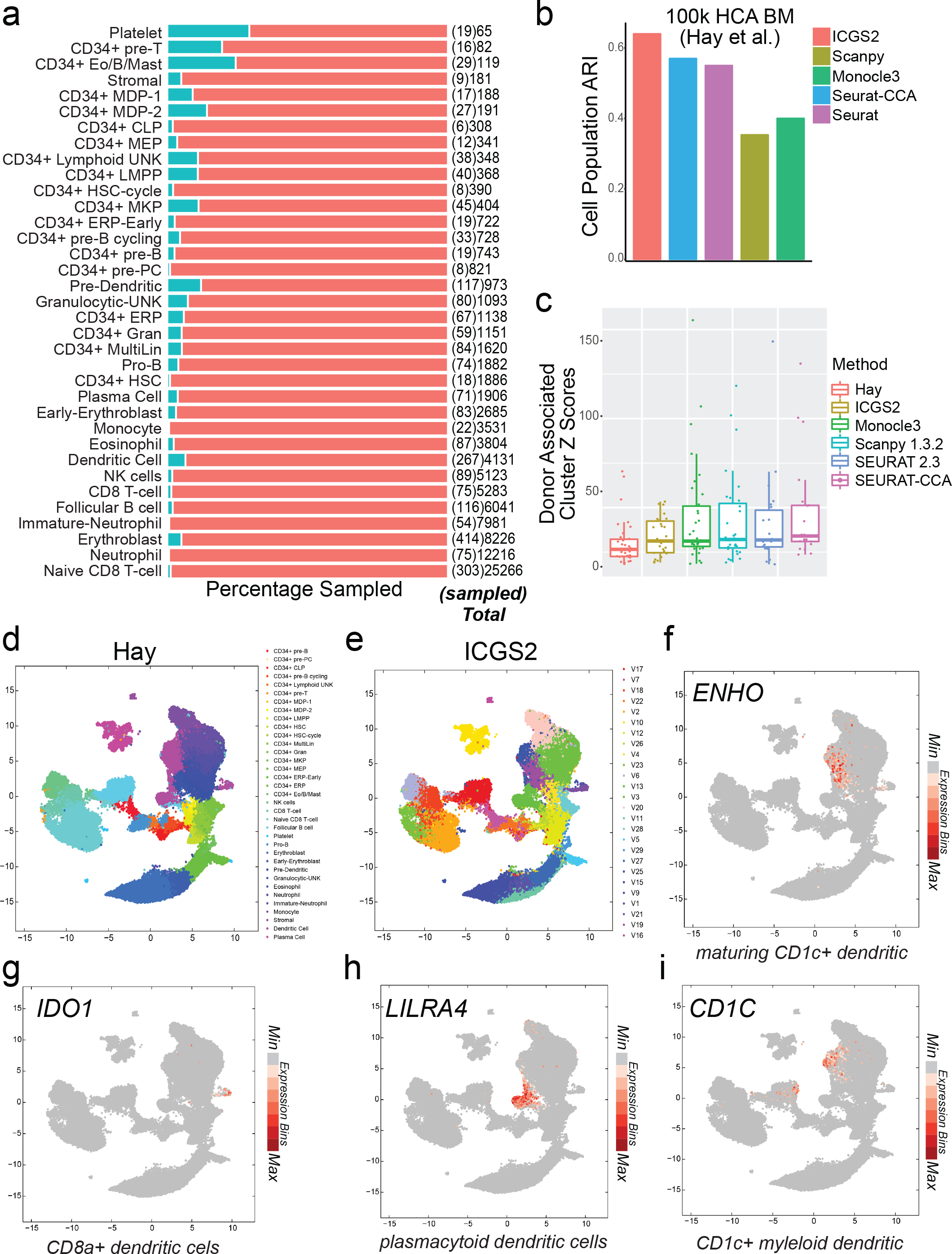
Identification of rare and novel cell populations from ultra large scRNA-Seq data. a) The frequency of PageRank down-sampled Bone Marrow cells from the Human Cell Atlas (HCA) project (n=100,514) are shown relative to the total number of cell prior to sampling for the prior defined cell-populations (n=35). The total number of cells in each cell population are shown to the right of the plot with the number of sampled samples in brackets. b) Comparison of prior-defined Bone Marrow clusters (ARI scores) using ICGS2-DS (down-sampled), Seurat v2.3, Seurat CCA, Monocle 3 and Scanpy. c) Comparison of the different algorithms in detecting donor biased Bone Marrow clusters (aka batch effects). Z-enrichment scores are calculated for each of the eight Bone Marrow donors against each cell cluster identified by the evaluated algorithm. A high z-score indicates an enrichment in cells in a specific cluster and a specific donor. Clusters enriched in cells from individual Bone Marrow donors indicated by high z-scores are visualized as outliers (circles) in the box-plots. d-e) UMAP visualization of clusters for prior-defined Bone Marrow HCA scRNA-Seq clusters by Hay et al. and by ICGS2 with down-sampling. f-i) 4 ICGS2 identified novel dendritic cell populations visualized using the top ranked ICGS2 gene marker.

## DISCUSSION

As scRNA-Seq approaches continue to increase in the depth of cells captured and molecules measured, more accurate approaches are required to identify rare and subtly distinct cell populations associated unique gene expression programs. Here we present an improved and highly scalable version of ICGS, that can be applied to extremely large scRNA-Seq datasets to delineate subtly distinct and rare cell populations. We use a hybrid approach that combines accurate methods for cluster determination and cell classification, in combination with new algorithms for intelligent single-cell down-sampling. NMF has been shown to improve the detection of sub-populations from diverse datasets, due to its ability to identify interpretable parts from high dimensional datasets ^32^. Using this refined workflow, we demonstrate improved performance over a large spectrum of existing approaches, across different datasets of varying complexity and size. Importantly, the use of iterative gene correlation and guide-gene selection appears to significantly minimize the impact of donor effects in the context of the first HCA release of human transcriptome variation, without directly considering such effects. This approach further uniquely identifies additional cell populations in bone marrow that decompose multiple prior defined cell-types associated with biologically informative markers (Multi-Lin, T-cells, Erythroblasts and Dendritic cells).

ICGS2 is fundamentally distinct from alternative approaches, in terms of its basic strategy to identify heterogeneity. Standard methods for variable gene selection (dispersion, PCA) are inherently more susceptible to initial transcriptional noise, batch and donor effects, however, ICGS selects variable genes through a rigorous pairwise correlation strategy over multiple rounds of iteration, with a focus on the selection of transcription factors as guide-genes. As previously demonstrated, this approach is more likely to identify transitional states which include mixed-lineage progenitors, weekly defined by unique gene expression ^6, 13, 21, 33, 34, 35^. ICGS2 extends the ability of ICGS to further define rare and common transcriptionally distinct populations, including multi-lineage cell populations from the human cell atlas, independent of donor effects. Because the software automatically identifies the most appropriate number of clusters, it can be simultaneously applied to many datasets, without the requirement for the user to specify.

The potential applications of this approach are broad, which include emerging large-scale whole-organism atlases, where AltAnalyze provides additional advantages beyond the ICGS2 algorithm itself. These benefits include imbedded methods to predict cell-type identify based on existing cell-specific gene-set references (gene set enrichment, cellHarmony), pathway enrichment analysis and display of protein-protein and transcriptional regulatory network relationships among genes differentially expressed between similar populations (NetPerspective algorithm) ^13^. Importantly, this workflow is accessible by both knowledgeable single-cell data analysts as well as conventional biologists without such expertise, through accessible command-line and graphical user interfaces. An important caveat of this approach is that it is dependent on the presence of coordinate gene expression patterns in which the underlying data is not so sparse that initially correlated genes can be identified. To address this challenge, this tool further includes the ability for users to designate the number of clusters when initial heterogeneity is only weakly detected. While the parameters of ICGS2 and other methods (e.g., SC3, Seurat) can be modified to identify additional subtypes, in the future, we hope to optimize our approach to optionally find maximal heterogeneity at the lowest resolution (sub-clustering). Through similar uses of ICGS2, we anticipate the discovery novel biologically informative cell populations that can guide our understanding of cellular diversity on complex organisms, including exceedingly rare populations that underlie disease phenotypes.

## References

1. Andrews TS, Hemberg M. Identifying cell populations with scRNASeq. Molecular Aspects of Medicine 59, 114–122 (2018).

2. Maaten Lvd, Hinton G. Visualizing Data using t-SNE. Journal of Machine Learning Research 9, 2579–2605 (2008).

3. McInnes L, Healy J. UMAP: Uniform Manifold Approximation and Projection for Dimension Reduction. arXiv:180203426 [cs, stat], (2018).

4. Kiselev VY, et al. SC3: consensus clustering of single-cell RNA-seq data. Nature Methods 14, 483–486 (2017).

5. Cao J, et al. The single-cell transcriptional landscape of mammalian organogenesis. Nature 566, 496 (2019).

6. Olsson A, et al. Single-cell analysis of mixed-lineage states leading to a binary cell fate choice. Nature 537, 698–702 (2016).

7. van der Laan MJ, Pollard KS. A new algorithm for hybrid hierarchical clustering with visualization and the bootstrap. Journal of Statistical Planning and Inference 117, 275–303 (2003).

8. Kim H, Park H. Sparse non-negative matrix factorizations via alternating non-negativity-constrained least squares for microarray data analysis. Bioinformatics 23, 1495–1502 (2007).

9. Cortes C, Vapnik V. Support-vector networks. Mach Learn 20, 273–297 (1995).

10. Emig D, Salomonis N, Baumbach J, Lengauer T, Conklin BR, Albrecht M. AltAnalyze and DomainGraph: analyzing and visualizing exon expression data. Nucleic acids research 38, W755–762 (2010).

11. Aumüller M, Bernhardsson E, Faithfull A. ANN-Benchmarks: A Benchmarking Tool for Approximate Nearest Neighbor Algorithms. arXiv:180705614 [cs], (2018).

12. Hubert L, Arabie P. Comparing partitions. Journal of Classification 2, 193–218 (1985).

13. Lu Y-C, et al. The Molecular Signature of Megakaryocyte-Erythroid Progenitors Reveals a Role for the Cell Cycle in Fate Specification. Cell Reports 25, 2083–2093.e2084 (2018).

14. Zeisel A, et al. Brain structure. Cell types in the mouse cortex and hippocampus revealed by single-cell RNA-seq. Science 347, 1138–1142 (2015).

15. Pollen AA, et al. Low-coverage single-cell mRNA sequencing reveals cellular heterogeneity and activated signaling pathways in developing cerebral cortex. Nature Biotechnology 32, 1053–1058 (2014).

16. Usoskin D, et al. Unbiased classification of sensory neuron types by large-scale single-cell RNA sequencing. Nat Neurosci 18, 145–153 (2015).

17. Treutlein B, et al. Reconstructing lineage hierarchies of the distal lung epithelium using single-cell RNA-seq. Nature 509, 371–375 (2014).

18. Guo M, Wang H, Potter SS, Whitsett JA, Xu Y. SINCERA: A Pipeline for Single-Cell RNA-Seq Profiling Analysis. PLOS Computational Biology 11, e1004575 (2015).

19. Xu C, Su Z. Identification of cell types from single-cell transcriptomes using a novel clustering method. Bioinformatics 31, 1974–1980 (2015).

20. Butler A, Hoffman P, Smibert P, Papalexi E, Satija R. Integrating single-cell transcriptomic data across different conditions, technologies, and species. Nature Biotechnology 36, 411–420 (2018).

21. Hay S, Ferchen K, Chetal K, Grimes HL, Salomonis N. The Human Cell Atlas bone marrow single-cell interactive web portal. Exp Hematol, (2018).

22. Group HCAW. HCA Data Coordination Platform. (ed^(eds) (2018).

23. Wolf FA, Angerer P, Theis FJ. SCANPY: large-scale single-cell gene expression data analysis. Genome Biol 19, 15 (2018).

24. Qiu X, et al. Reversed graph embedding resolves complex single-cell trajectories. Nature Methods 14, 979–982 (2017).

25. Iacono G, et al. bigSCale: an analytical framework for big-scale single-cell data. Genome Res, (2018).

26. Page L, Brin S, Motwani R, Winograd T. The PageRank Citation Ranking: Bringing Order to the Web (1998).

27. Heger L, et al. CLEC10A Is a Specific Marker for Human CD1c+ Dendritic Cells and Enhances Their Toll-Like Receptor 7/8-Induced Cytokine Secretion. Front Immunol 9, 744–744 (2018).

28. Eggink LL, Roby KF, Cote R, Kenneth Hoober J. An innovative immunotherapeutic strategy for ovarian cancer: CLEC10A and glycomimetic peptides. J Immunother Cancer 6, (2018).

29. Yan Z, et al. A novel peptide targeting Clec9a on dendritic cell for cancer immunotherapy. Oncotarget 7, 40437–40450 (2016).

30. Orabona C, et al. Toward the identification of a tolerogenic signature in IDO-competent dendritic cells. Blood 107, 2846–2854 (2006).

31. Cao W, et al. Plasmacytoid dendritic cell-specific receptor ILT7-Fc epsilonRI gamma inhibits Toll-like receptor-induced interferon production. J Exp Med 203, 1399–1405 (2006).

32. Mejía-Roa E, Tabas-Madrid D, Setoain J, García C, Tirado F, Pascual-Montano A. NMF-mGPU: non-negative matrix factorization on multi-GPU systems. BMC Bioinformatics 16, 43 (2015).

33. Hulin A, et al. Maturation of heart valve cell populations during postnatal remodeling. Development, (2019).

34. Magella B, et al. Cross-platform single cell analysis of kidney development shows stromal cells express Gdnf. Dev Biol 434, 36–47 (2018).

35. Yáñez A, et al. Granulocyte-Monocyte Progenitors and Monocyte-Dendritic Cell Progenitors Independently Produce Functionally Distinct Monocytes. Immunity 47, 890–902.e894 (2017).

